# Evolution leads to emergence: An analysis of protein interactomes across the tree of life

**DOI:** 10.1101/2020.05.03.074419

**Authors:** Erik Hoel, Brennan Klein, Anshuman Swain, Ross Grebenow, Michael Levin

## Abstract

The internal workings of biological systems are notoriously difficult to understand. Due to the prevalence of noise and degeneracy in evolved systems, in many cases the workings of everything from gene regulatory networks to protein-protein interactome networks remain black boxes. One consequence of this black-box nature is that it is unclear at which scale to analyze biological systems to best understand their function. We analyzed the protein interactomes of over 1800 species, containing in total 8,782,166 protein-protein interactions, at different scales. We demonstrate the emergence of higher order ‘macroscales’ in these interactomes and that these biological macroscales are associated with lower noise and degeneracy and therefore lower uncertainty. Moreover, the nodes in the interactomes that make up the macroscale are more resilient compared to nodes that do not participate in the macroscale. These effects are more pronounced in interactomes of Eukaryota, as compared to Prokaryota. This points to plausible evolutionary adaptation for macroscales: biological networks evolve informative macroscales to gain benefits of both being uncertain at lower scales to boost their resilience, and also being ‘certain’ at higher scales to increase their effectiveness at information transmission. Our work explains some of the difficulty in understanding the workings of biological networks, since they are often most informative at a hidden higher scale, and demonstrates the tools to make these informative higher scales explicit.

## Introduction

Interactions in biological systems are noisy and degenerate in their functions, making them fundamentally noisier and fundamentally different from those in engineered systems (Edelman & Gally, 2001; Tsimring, 2014). The sources of noise in biology are nearly ubiquitous and vary widely. Noise may exist a gene regulatory network, wherein a gene might upregulate another gene but only probabilistically, or they may be noisy in that a protein may bind randomly across a set of possible pairings. There are numerous sources of such *indeterminism* in cells and tissues, such as how cell molecules a buffered by Brownian motion (Einstein, 1905), to the stochastic opening and closing of ion channels (Colquhoun & Hawkes, 1981), and even to the chaotic dynamics of neural activity (Başar, 2012).

There are also numerous sources of degeneracy within the cellular, developmental, and genetic operation of organisms (Brennan, Cheong, & Levchenko, 2012). Degeneracy is when an end state or output, like a phenotype, can come from a large number of possible states or inputs networks (Tononi, Sporns, & Edelman, 1999).

Due to this indeterminism and degeneracy, the dynamics and function of biological systems are often uncertain. This hampers control of system-level properties for biomedicine and synthetic bioengineering, as well as hampering the understanding of modelers and experimentalists who wish to build “big data” approaches to biology like interactomes, connectomes, and mapping molecular pathways (Dolinski & Troyanskaya, 2015; Marx, 2013). While there have been many attempts to characterize and understand this uncertainty in biological systems (Tononi, Sporns, & Edelman, 1999), the explanations typically do not extend beyond the advantages of redundancy in these systems (Whitacre, 2010).

How do noise and uncertainty span the tree of life? Here we examine this question in biological networks, a common type of model for biological systems networks (Alon, 2003; Bray, 2003). Specifically, we examine protein-protein interactomes from organisms across a wide range of organisms to investigate whether or not the noise and uncertainty in biological networks increases or decreases across evolution. In order to quantify this noise and uncertainty we make use of the *effective information* (*EI*), an information-theoretic network quantity based on the entropy of random walker behavior on a network. A lower *EI* indicates greater noise and uncertainty (a formal mathematical definition is given in the Results). Indeed, the *EI* of biological networks has already been shown to be lower in biological networks compared to technological networks (Klein & Hoel, 2020), which opens the question of why this is the case.

To see how *EI* changes across evolution, we examined networks of protein-protein interactions (PPIs) from organisms across the tree of life. The dataset consists of *interactomes* from 1840 species (1,539 Bacteria, 111 Archaea, and 190 Eukaryota) derived from the STRING database (Szklarczyk et al., 2010; Szklarczyk et al., 2017). These interactomes have been previously used to study evolution of resilience, where researchers found that species tended to have higher values of *network resilience* with increasing evolution (wherein “evolution” was defined as the number of nucleotide substitutions per site) (Zitnick et al. 2019). In our work, we take a similar approach, highlighting changes in interactome properties as evolution progresses.

Additionally, we focus on identifying when interactomes have *informative macroscales*. A *macroscale* refers to some dimension reduction, such as an aggregation, coarse-graining, or grouping, of states or elements of the biological system. In networks this takes the form of replacing subgraphs of the network with individual nodes (*macro-nodes*). A network has an *informative* macroscale when subgraphs of the network can be grouped into macro-nodes such that the resulting dimensionally-reduced network gains *EI* (Klein & Hoel, 2019). When such grouping leads to an increase in *EI*, we describe the resulting macro-node as being part of an informative macroscale. Following previous work, we refer to any gain in *EI* at the macroscale as *causal emergence* (Hoel et al., 2013). With these techniques, we can identify which protein-protein interaction (PPI) networks have informative macroscales and which do not. By correlating this property with where (in time) each species lies in the evolutionary tree, we show that informative macroscales tend to emerge *later* in evolution, being associated more with Eukaryota than Prokaryota (such as Bacteria).

What is the evolutionary advantage of having informative higher scales? This question is important because higher scales minimize noise or uncertainty in biological networks. Yet such uncertainty or noise represents a fundamental paradox. The more noisy a network is, the more uncertain and the less effective that network is (like being able to effectively transform inputs to outputs, such as being able to effectively upregulate a particular gene in response to a detected chemical in the environment). Therefore, we might expect evolved networks to be highly effective. Yet this is the opposite of what we observe. Instead we observe that effectiveness of lower scales decreases later in evolution, as higher scales that are effective emerge.

We argue here that this multi-scale behavior is the resolution to a paradox: there are advantages to being effective, but there are also advantages to being less effective and therefore more uncertain or noisy. For instance, less effective networks might be more resistant to attack or node failure due to redundancy. The paradox is that networks that are certain are effective yet are vulnerable to attacks or node failures, while networks that are uncertain are less effective but are resilient in the face of attacks or node failures. We argue that biological networks have evolved to resolve this “certainty paradox” by having informative higher scales. Specifically, we propose that the macroscales of a biological network evolve to have high effectiveness, but their underlying microscales may have low effectiveness, therefore making the system resilient without paying the price of a low effectiveness.

In a biological sense, node failures or attacks in a cellular network may represent certain mutations in proteins or other biochemical entities, which in turn may prevent regular functioning of the system (Barabási & Oltvai, 2004). Biological networks should then, over the course of evolution, develop degeneracy and noise at lower scales to maintain regular functioning, while at the same time developing effectiveness at a higher level. This transformation can be achieved by the action of both neutral and selective processes in evolution. Neutral processes such as pre-suppression, which aided by mutations, increases the number of interactions (Lukeš et al., 2011) and can therefore decrease network effectiveness. On the other hand, selective processes can weed out the noise that interferes with the functioning and efficiency of the system (Brunet & Doolittle, 2018). An interplay of these evolutionary processes can lead to a resolution the “certainty paradox” in cellular networks by the develop of informative macroscales.

This work therefore presents an explanation for the observed trend in increased resiliency through evolution (Zitnick et al., 2019): informative macroscales make networks more resilient. Finally, we offer insights into biological processes at molecular level that might be responsible for the emergence of informative macroscales in protein-protein interaction networks, specifically looking at the differences between Bacteria, which has a low rate of nucleotide substitutions per site, and Eukaryota, which exhibit a higher rate. Understanding the basic principles governing the differences in efficiency and uncertainty between these major divisions of life can help us comprehend the trade-offs involved in information processes in PPIs across evolution.

## Results

### Effectiveness of protein interactomes across the tree of life

Effective information (*EI*) is a network property reflecting the certainty (or uncertainty) contained in that network’s connectivity (Klein & Hoel, 2020). It is a structural property of a network calculated by traversing its topology, and is based on the uncertainty of a random walker’s transitions between pairs of nodes, and the distribution of this uncertainty throughout the network. It is calculated by examining the network’s connectivity.

In a protein interactome, the nodes are individual proteins and the edges of the network are interactions, generally describing the possibility of binding between two proteins. Therefore, the uncertainty we analyze is uncertainty as to which protein(s) a given protein might interact with, e.g., bind with. Each node in the network has out-weights, which are represented by a vector *W*_*i*_^*out*^. For instance, protein *A* might share an edge with protein *B* and also protein *C*. Therefore, *W*_*A*_^out^. is [½, ½]. Since most protein interactome are undirected, its edges are normalized for each node (such that the sum of *W*_*i*_^*out*^ for each node is 1.0). Note that this process of normalization implies that the probability of binding is uniform across the different possible interactions. This transformation into a direct network makes the networks amenable to standard tools of network science, such as analyzing random walk dynamics, and it is also necessary to calculate the *EI* of the network. Additionally, the uniform distribution of 1/*n* is the simplest a priori assumption. However, the actual probability of binding is dependent on biological background conditions such as protein prevalence and not included in most open-source models, and therefore our analysis could change if such detailed probabilities were known.

The uncertainty associated with each protein can be mathematically captured by examining the entropy of the outputs of a node, *H*(*W*_*i*_^*out*^), wherein a higher entropy indicates more uncertainty more uncertainty as to interactions (Shannon, 1948). The entropy of the distribution of weight across the entire network, *H*(〈*W*_*i*_^*out*^〉), reflects the spread of uncertainty across the network. A lower *H*(〈*W*_*i*_^*out*^〉) means that information is distributed only over a small number of nodes. A high *H*(〈*W*_*i*_^*out*^〉) signifies that information is dispersed throughout the network. The *EI* of a network can then be defined as the entropy of distribution of weights over the network minus the average uncertainty inherent in the weight of each node, or:

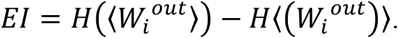

*EI* can itself be further decomposed into the degeneracy and indeterminism of a network (Klein & Hoel, 2020), where each indicate the lack of specificity in the network’s connectivity or interactions. Degeneracy indicates a lack of specificity in targeting nodes (many nodes target the same node), while indeterminism indicates a lack of specificity in targeted nodes (nodes target many nodes). Note that, if networks are considered deterministic in the physical sense, the indeterminism term of *EI* still reflects the uniqueness of targets in the network.

A network where all the nodes target a single node will have zero *EI* (since it has maximum degeneracy), as will a network where all nodes target all other nodes (complete indeterminism). *EI* will only be maximal if every node has a unique output. This forces the *EI* of a network to be bounded by log_2_(*n*). Therefore, in order to compare networks of different sizes, *EI* can be normalized using the number of nodes in the network, *n*, and a new quantity, *effectiveness*, can be defined as:

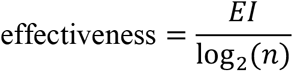

To explore the change in effectiveness of biological networks, we examined protein interactomes of 1840 species divided between Archaea, Bacteria, and Eukaryota (see the Methods section for details on the origin and nature of these protein interactomes). We found a clear pattern in the effectiveness of the networks, based on where they are located in the tree of life (Fig. 1), the position of which is based on each protein interactome’s small subunit ribosomal RNA gene sequence information (Hug et al., 2016) (see Methods for details). Overall, we found that the mean effectiveness of protein interactomes actually decreases later in the tree of life as nucleotide substitutions occurred. Specifically, Bacteria were found to have a greater effectiveness (0.77) compared to Eukaryota (0.72) on average (student’s t-test, *p* < 10^−8^). Following Zitnik et al. (2019), we restricted further statistical analysis to interactomes with more than 1000 citations, in order use the most well-founded protein interactomes, but the directionality and significance of the result is unchanged when only those above 100 citations are included as compared to when all interactomes are included (student’s t-test, *p* < 10^−11^). Due to the small number of Archaea interactomes based on above 1000 citations we did not include those samples in Fig. 1B.

**Figure 1.**
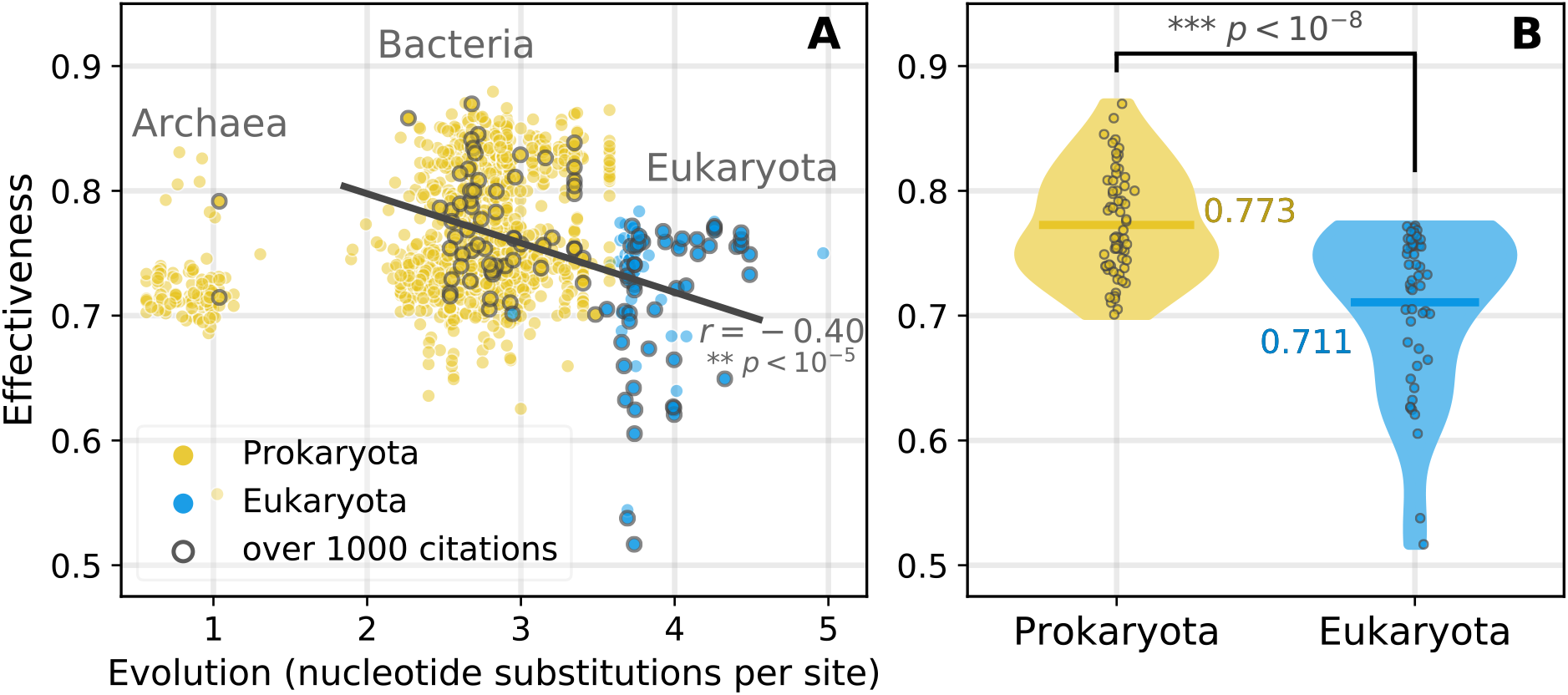
Effectiveness of protein interactomes. **(A)** Effectiveness of all 1840 species with their superphylum association. Interactomes with a lower number of nucleotide substitutions per site tended to be Prokaryota (yellow), while those higher tended to be Eukaryota (blue). Solid line is a linear regression comparing the effectiveness of Bacteria and Eukaryota (*r* = −0.40, *p* < 10^−5^), due to the small number of Archaea that passed the threshold for reliable datasets (see Results). **(B)** The effectiveness of prokaryotic protein interactomes is greater than that of eukaryotic species, indicating that effectiveness might decrease with more nucleotide substitutions per site.

### Causal emergence across the tree of life

At first the higher effectiveness in Prokaryota interactomes as compared to that of Eukaryota (as shown in Fig. 1) may seem counter-intuitive. One might naively expect the effectiveness of cellular machinery, including or especially interactomes, to increase over evolutionary time, instead of decreasing as we have shown.

One hypothesis to explain these results is that, while the protein interactomes get less effective in their micro-scales over evolutionary time, the interactomes are able to nonetheless be effective due to the emergence of informative macroscales as evolution proceeds. To examine this hypothesis, we must first define a procedure for finding macroscales in networks.

Network macroscales are defined as subgraphs (i.e., connected sets of nodes and their associated links) that can be grouped into single macro-nodes such that the resulting network has a higher value of *EI* than the original microscale network (Klein & Hoel, 2020). We denote the microscale of a network as *G* and the macroscale as *G*_*M*_, which is composed of both ungrouped nodes (micro-nodes) and macro-nodes, μ. The macroscale network, *G*_*M*_, is a dimension reduction in that it always has fewer nodes than *G*.

To recast a particular subgraph into a macro-node, its connectivity must be modified since the subgraph is being transformed into a single node. In terms of input to the new macro-node, μ, all out-weights that targeted nodes in the subgraph now target the macro-node. In terms of output, each micro-node, *v*_*i*_, inside the subgraph has some *W*_*i*_^*out*^. To recast the nodes inside a subgraph into a macro-node, we replace the *W*_*i*_^*out*^ of the nodes in the subgraph a single *W*_μ_^*out*^, which is a weighted average of the set of each *W*_*i*_^*out*^ in the subgraph. The weight is based off the probability *p* of each node *v*_*i*_ in the stationary distribution of the network, π. This forms macro-nodes (μ|π) that accurately recapitulate the microscale random walk dynamics at the new macroscale (Klein & Hoel, 2020). A macroscale is *informative* if it increases the *EI* of the network compared to the original microscale. In order to find macro-nodes that maximally increase *EI* we make use of a modified spectral algorithm to find locally optimal micro-to-macro mappings, originally described in (Griebenow et al., 2019).

Results from this analysis support our initial hypothesis that effectiveness is actually being transitioned to macroscales of biological networks in Eukaryota over evolutionary time, even though the microscales become noisy and less effective over evolutionary time. The total amount of *causal emergence* (the gain of *EI* by grouping subgraphs into macro-nodes) was identified for each protein interactome from each species, normalized by the total size of that protein interactome (Fig 2A). Across the tree of life we observe that Eukaryota have more informative macroscales and show a significant difference in the percentage of microscale nodes that get grouped into macro-nodes than (Fig. 2).

**Figure 2.**
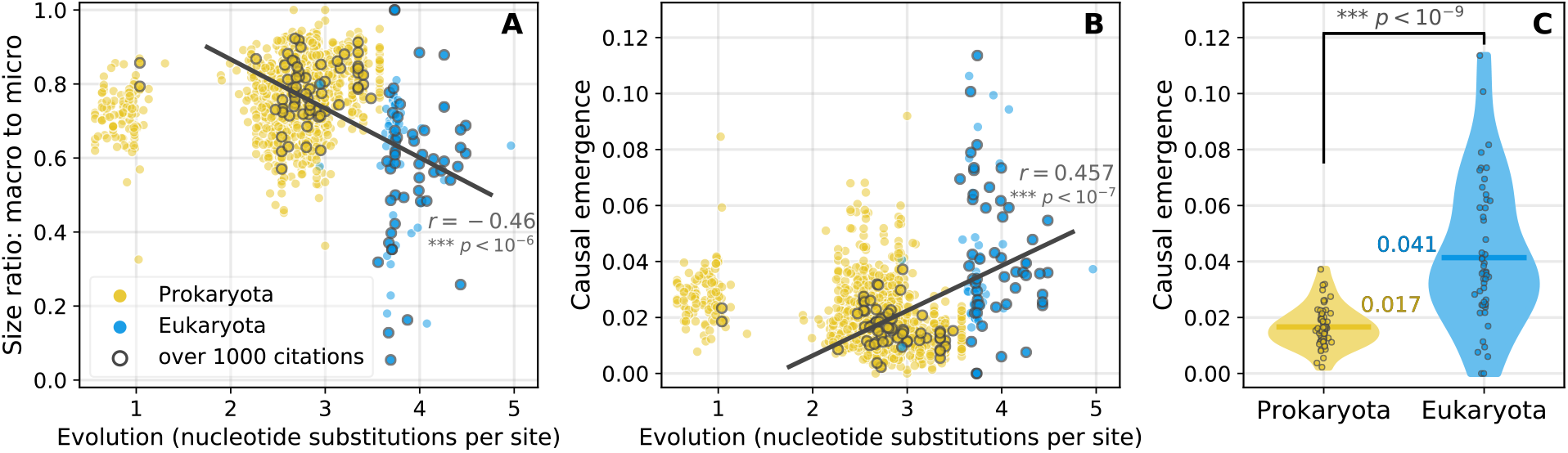
Causal emergence in protein interactomes. **(A)** The protein interactomes of each species undergoes a modified spectral analysis in order to identify the scale with *EI*^*max*^. The total dimension reduction of the network is shown, with there being a greater effect in Eukaryota as more subgraphs are grouped in macro-nodes. That is, as evolutionary time goes on the coarse-grained networks become a smaller fraction of their original microscale network size (*r* = −0.46, *p* < 10^−6^). **(B)** In order to compare the degree of causal emergence in protein interactomes of different sizes, the total amount of causal emergence is normalized by the size of the network, log_2_(*n*), and we see here a positive correlation between evolution and causal emergence (*r* = 0.457, *p* < 10^−7^). **(C)** The amount of normalized causal emergence is significantly higher for Eukaryota.

### Macroscales of protein interactomes are more resilient than their microscales

Why might biological networks evolve over time to have informative macroscales? As previously discussed, one answer might be that having multi-scale structure provides benefits that networks with only a single scale lack. All networks face a “certainty paradox.” The paradox is that uncertainty in connectivity is desirable since it is protective from node failures. For instance, a node failure could be the removal of a protein due to a nonsense mutation, or the inability to express a certain protein due to an environmental effect, such as a lack of resources, or even a viral attack. In turn, this could lead to a loss of biological function or the development of disease or even cell death. A protein interactome may be resilient to such node failures by being highly uncertain or degenerate in its protein-protein interactomes. However, this comes at a cost. A high uncertainty can lead to problems with reliability, uniqueness, and control in terms of effects, such as an inability for a particular protein to deterministically bind with another protein. For instance, in a time of environmental restriction of resources, certain protein-protein interactomes may be necessary for continued cellular function, but if there is large-scale uncertainty even significant upregulation of genes controlling expression may not lead reliably to a certain interaction.

Here we explore these issues by examining the *network resilience* of protein interactomes in response to node removals, which represent either attacks or general node failures.

In order to measure the resilience of the network in response to a node removal we follow (Zitnick et al., 2019) by using the change in the Shannon entropy of the component size distribution of the network following random node removal. That is, if *p*_*c*_ is the probability that a randomly selected node is in connected component *c* ∈ *C* following the removal of a fraction *f* of the nodes in the network, the entropy associated with the component size distribution, *H*(*G*_*f*_), is:

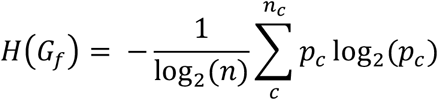

where *n*_*c*_ is the number of connected components remaining after *f* fraction nodes have been removed (note: “removed” here indicates that the nodes become isolates, still contributing to the component size distribution though not retaining any of the original links). The change in entropy,

*H*(*G*_*f*_), as *f* increases from 0.0 to 1.0 corresponds to the resilience of the network in question. Specifically, this resilience is defined as follows:

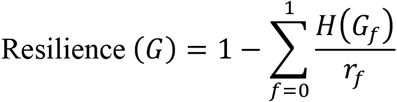

where *r*_*f*_ is the rate of node removal (i.e., the increment that the fraction *f* increases from 0.0 to 1.0). In this work, we default to a value of *r*_*f*_ = 100, which means that the calculation of a network’s resilience involves iteratively removing 1%, 2%, … 100% of the nodes in the network. For each value of *f*, we simulate the node removal process 20 times.

Our hypothesis is that biological networks deal with this “certainty paradox” by maintaining uncertainty at their microscale. This gives a pool of noise and degeneracy, leading to resilience. Meanwhile, at the macroscale, the networks can develop a high effectiveness, wherein sets of proteins deterministically and non-degenerately interact. To explore this hypothesis, we compare the network’s resilience to removing micro-nodes that are members of subgraphs grouped into macro-nodes to the network’s resilience to removing micro-nodes that remain ungrouped (shown in Fig. 3).

**Figure 3.**
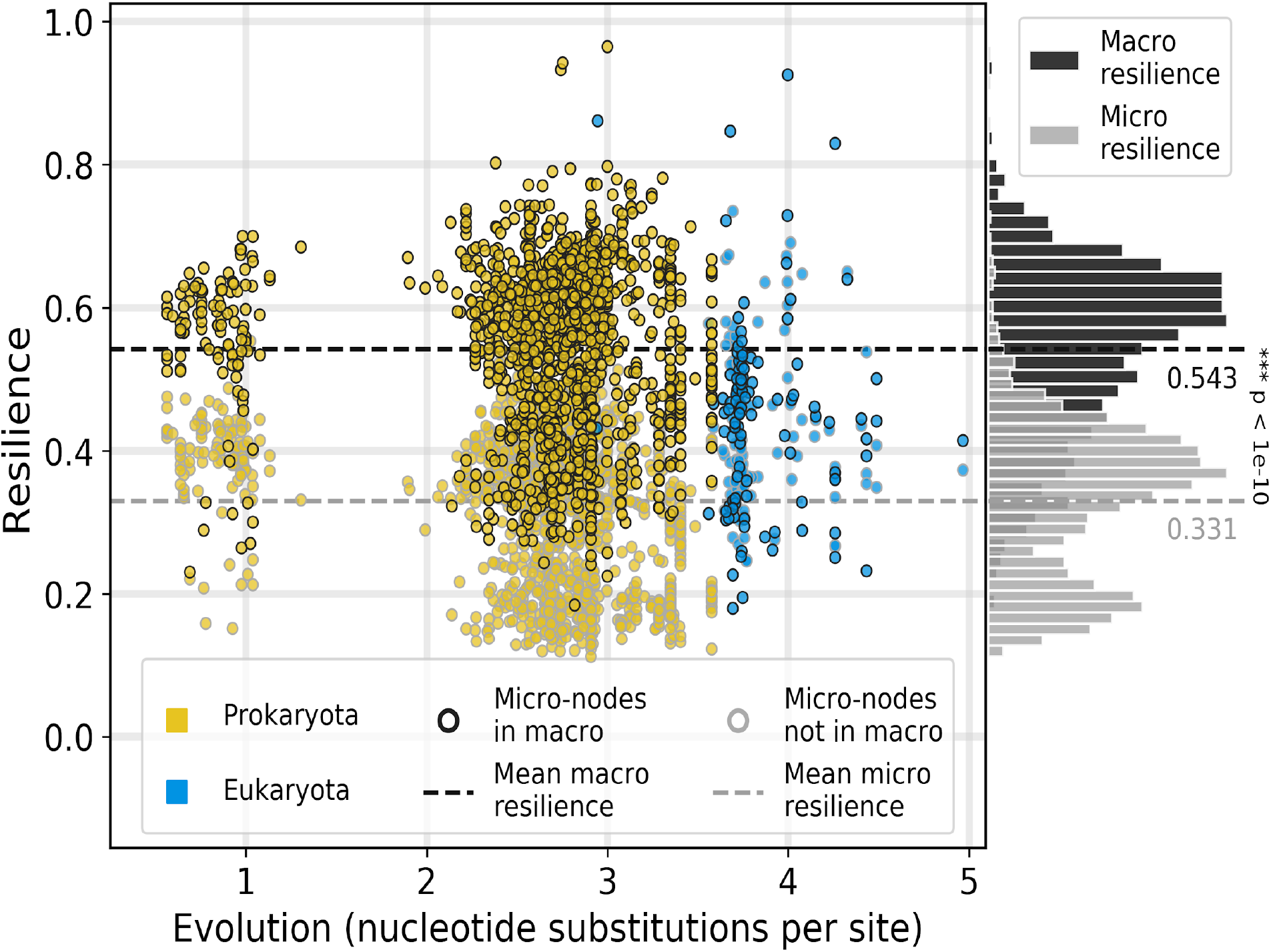
Resilience of micro- and macro-nodes following causal emergence in interactomes. The resilience of a species interactome changes across the tree of life, as shown in previous research (Zitnick et al., 2019). Using the mapping generated by computing causal emergence (Fig. 2B), we calculate the resilience of the network, isolating the calculation to nodes that are either part of the macroscale or microscale. Points are color-coded according to the evolutionary domain; points with dark outlines are associated with micro-nodes that have been grouped into a macro-node (macroscale), while the points with light outlines have not been grouped into a macro-nodes (microscale). Nodes at the microscale contribute less to the overall resilience of a given network (0.331) compared to nodes that contribute to macro-nodes (0.543) on average (t-test, *p* < 1.0^−10^). Note: plotted are the microscale and macroscale resilience values for each interactome in the dataset, and the difference in resilience across scales holds even when only including species with more than 10, 100, or 1000 citations.

By isolating the calculation of network resilience to only the micro- or macro-nodes of a network, we see a stark trend emerge wherein nodes inside highly informative macro-nodes are more resilient than nodes outside. That is, nodes in the original interactome that were grouped into a macro-node contribute more to the overall resilience of the interactome. This not only supports our hypothesis that biological networks resolve the “certainty paradox” by building multi-scale structure, but also provides further explanation and contextualization for the recent findings of increasing resilience across evolutionary time (Zitnick et al., 2019).

## Discussion

In this work, we analyzed how the informativeness of protein interactomes changed over evolutionary time. Specifically, we made use of the *effective information* (*EI*) to analyze the amount of uncertainty (or noise) in the connectivity of protein interactomes. We found that the effectiveness (the normalized *EI*) of protein interactomes decreased over evolutionary time, indicating that uncertainty in the connectivity of the interactomes was increasing over evolutionary time. However, we discovered that this was due to eukaryotic protein interactomes possessing higher (informative) scales, such that they had more *EI* when recast as a coarse-grained network— a phenomenon known as causal emergence. This lower effectiveness and higher causal emergence in eukaryotic species was due to the indeterminism and degeneracy in the network structure of their protein-protein interactions.

We used a dataset from the STRING database (Szklarczyk et al., 2010; Szklarczyk et al., 2017) that spans more than 1800 species (1,539 Bacteria, 111 Archaea, and 190 Eukaryota), which has been shown to have considerable advantages compared to previous collections of protein interactomes (Zitnik et al., 2019). However, we cannot rule out the possibility that biases might exist in the specific manner of data collection, such as high under-representation of specific types of difficult-to-detect interactions, which could potentially introduce errors in the calculations of effectiveness in eukaryotic interactomes. As such, we conducted a series of statistical robustness tests that accounted for potential biases in both the data collection and network structures of interactomes in our dataset (see Fig. 4 in Methods for further details about these statistical tests). In short, the results we observed in this study cannot be explained by two plausible sources of bias: 1) Random rewiring of network edges does not produce similar results and 2) Network null models of each interactome in this study produce only a fraction of the observed causal emergence in our dataset (the maximum causal emergence values for a species’ network null model only reached 3% of the causal emergence of the original interactome). Notwithstanding these statistical tests, as technology and methods continue to improve these results and hypothesis must be tested rigorous.

**Figure 4.**
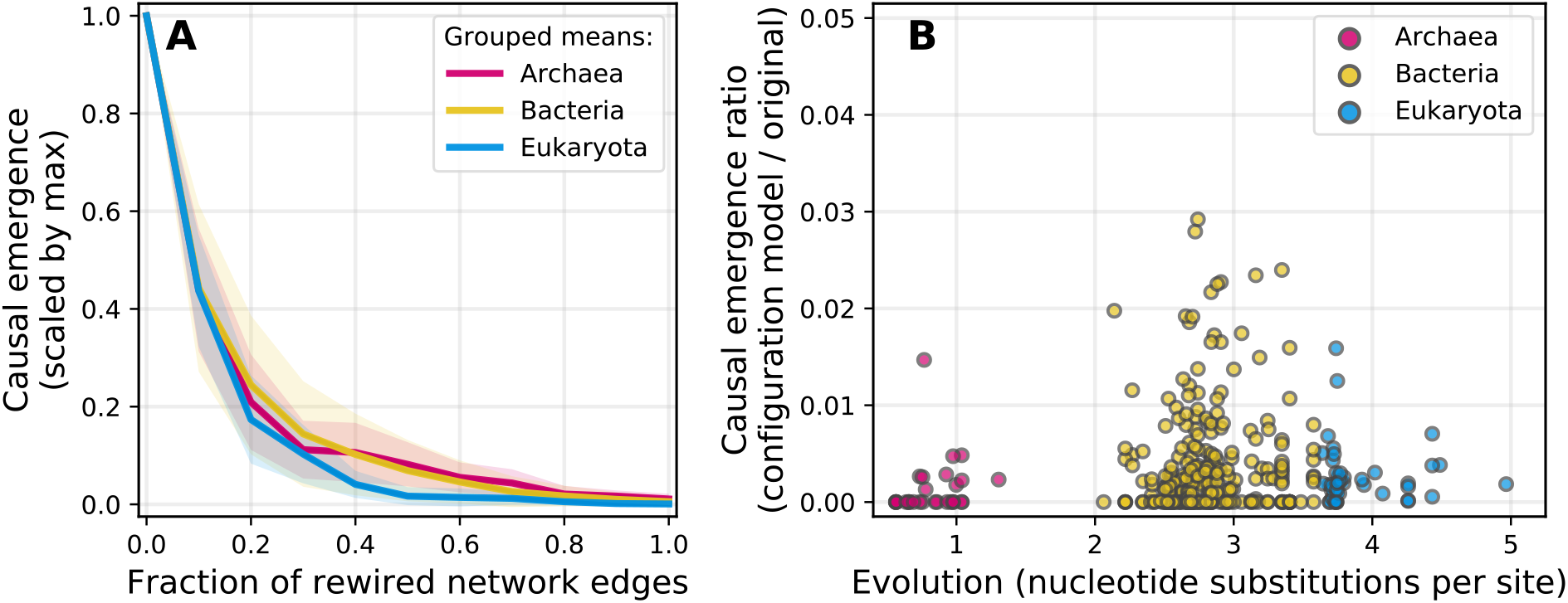
Statistical controls and network robustness tests. **(A)** As a greater fraction of network links are randomly rewired, we observe a decrease in the causal emergence values of the resulting networks (normalized by the causal emergence value of the original network). This appears not to be dependent on evolutionary domain, network size, density, or other network properties. **(B)** A second statistical control known as a *soft configuration model* assesses whether there is anything intrinsic to the network’s degree distribution that could be driving a given result. Here, we divide the average causal emergence of 10 such configuration model networks by the causal emergence values of the original protein interactome and observe that the null model networks preserve only a small fraction of the original amount of information gain (at most, the configuration models may show 3% of the original causal emergence).

To analyze why macroscales of biological networks evolved, we calculated how resilience differed for nodes inside of or outside of macro-nodes. We found that resilience of nodes left outside the macro was far lower, on average, than the resilience of nodes grouped into macro-nodes. This indicates that there are benefits of having macroscales, such as increased resilience, and that systems with informative macroscales can still have a high effectiveness but also maintain the benefits of having low effectiveness at a microscale. This is in line with the existing research showing that resilience increases with evolution (Zitnik et al., 2019).

These findings present evidence that biological systems are sensitive to the tradeoff between of effectiveness and robustness by examining whether evolution brings about multi-scale structure in biological networks. Systems with a single level of function face an irresolvable paradox: uncertainty in the connections and interactions between nodes leads to resilience to attack and robustness to node failures, but this decreases the effectiveness of that network. However, multi-scale systems, defined as those with an informative higher scale, can solve this “certainty paradox” by having high uncertainty in their connectivity at the microscale while having high certainty in their connectivity at the macroscale. The tradeoffs between being effective at a microscale (typically in prokaryotes, e.g. Bacteria) and being noisy at microscale, while transitioning the information to higher scales (Eukaryota) might have played a key role in evolutionary dynamics. Indeed, the drive from a prokaryotic ancestor to a eukaryotic one might have occurred based on this trade-off, however explaining such a phenomenon is outside the scope of the current work.

While we have illuminated many of the advantages of biological macroscales and posited a functional reason for their existence as the solution to the “certainty paradox,” what are the biological mechanisms behind the evolution of multi-scale structure? We offer here a few hypotheses about biological mechanisms that are concordant with the hypothesis of multi-scale advantages in terms of having both effectiveness and robustness.

Notably, evolution can proceed both via neutral processes and selection-based contexts. A well-known neutral process that affects interactions at cellular scale, such as those between proteins, is pre-suppression (also termed constructive neutralism) (Brunet & Doolittle, 2018). This refers to the complexity arising in the dependencies between interacting molecules in the absence of positive selection (Lukeš et al., 2011). Simply put, the likelihood of maintaining independence between partners is less than that of moving away from the original state (by accumulating changes), and therefore, random changes can increase the number of interactions between proteins in a system by chance alone, and result in “noisiness” in the interactions. This may offer a biological mechanism behind the result in low effectiveness in an interactome. Because Eukaryota have both larger number of proteins and a higher substitution rate than bacteria (Zitnik et al., 2019), eukaryotic interactomes might be expected to feature a higher number of neutral processes, all of which would combine to make interaction networks noisier and less effective. One hypothesis is that neutral evolution specifically drives the noise at the microscale but not the macroscale. At the macroscale interactomes would be trimmed and evolved under evolutionary constraints and selective pressures (Jain, Rivera, & Lake, 1999), which would eventually reinforce beneficial relationships, thinning out those that can cause negative effects on survival or growth (Brunet & Doolittle 2018). These processes may lead to formation of sub-groups of proteins in the network with more and stronger interactions *within* the group compared to fewer or weaker interactions between those in different subgroups (Brunet & Doolittle, 2018; Martin & Koonin, 2006)—thereby leading to the emergence of modular, macroscale structures in these networks, which we hypothesize to be correlated with organismal function (Alon, 2003).

Another possible explanation as to the biological mechanism behind our observed results of a decrease in effectiveness is that prokaryotes are more metabolically diverse than eukaryotes, possessing more metabolic processing pathways (Carlile, 1982). Together with changed usage patterns (such as carbon catabolite repression in Bacteria), this specificity of metabolite processing reduces energy demand and allows for more effective usage of resources (Gorke & Stulke, 2008). These processes would make biochemical inputs and outputs more streamlined and efficient in prokaryotes, which in turn, should increase the effectiveness of their protein interactomes, given energy and genomic size constraints (Giovannoni, Thrash, & Temperton, 2014). In contrast, Eukaryota, as a group, are less constrained by energy than prokaryotes (Lane, 2011) but must contend with a constrained number of metabolites, channelizing them to perform cellular functions in morphologically more complex environments (Carlile, 1982; Lane, 2011). Eukaryotic cells are about three orders of magnitude larger than prokaryotes (Lane, 2011), requiring more and different sets of controls and organizational processes. Prokaryotes depend on free diffusion for intracellular transport whereas Eukaryota have elaborate mechanisms for targeted transfers (Dacks, Peden, & Field; 2009). This reliance on cellular transport mechanisms can lead to higher modular (and thus more *degenerate* or *indeterministic*) structure in protein interactomes and other intracellular entities, which, as we show here, can be associated with *less* noise at higher scales of interaction. These higher-scale inter-module transfer mechanisms ensure the proper and less noisy flow of important molecules among these modules (such as protein or metabolite transport among organelles) (Alon, 2003). Each of these larger-scale processes, such as transport among organelles, relies on only a handful of inputs and outputs from outside its module, as compared to much more diverse interactions within the modules themselves (Alon, 2003), which arise due to both functional and neutral processes. In terms of networks, this hierarchical organizational structure is apt to lead to a higher network effectiveness score at the module/process scale compared to the micro-scale.

Such mechanistic biological explanations for why we might observe these differences in effectiveness are in line with the theoretical reasoning that biological systems need to resolve the paradox they face at individual scales and therefore construct multi-scale structure. We seek to tie the “certainty paradox” directly to the notion of *scale* in biological systems and provide a means for researchers to reduce the “black box” nature of these systems by searching across scales for models with low uncertainty. Understanding the mechanics of information transfer and noise in biological systems, and how they affect functionality, remains a major challenge in biology today. One can imagine that the drive from unicellular to multicellular life was based on some form of similar trade-offs, as those between prokaryotes and eukaryotes, that allowed multicellular life to operate via effective macro-states while reserving a pool of noise and degeneracy. Thus, understanding the information structure of these interactomes lends us an eye into the inner workings of long-term evolutionary processes and trade-offs that might have resulted in the two biggest phenotypic splits in evolutionary history—that of prokaryotic and eukaryotic cells, and of unicellular and multicellular life. We hope this developed framework is applied to other interactomes and other biological networks, such as GRNs, or even functional brain networks, to examine both how uncertainty plays a role in robustness, how informative higher scales change across evolution, and what fundamental tradeoffs biological systems face.

## Methods

### Protein interactomes

Protein interactomes are complex models of intracellular activity, often based on high-throughput experiments (Rual et al., 2005; Rolland et al., 2014). Here protein interactomes formed from a curated set of high-quality interactions between proteins (protein-to-protein interactions, or PPIs) are taken from the STRING database (Szklarczyk et al., 2010; Szklarczyk et al., 2017) the curation of which is outlined in (Zitnick et al., 2019). In this curation the STRING database (Search Tool for the Retrieval of Interacting Genes/Proteins, found at http://string-db.org) is used to derive a protein interactome for each species. Each PPIs in the protein interactome is an undirected edge where the edges are based on experimentally-documented physical interactions in the species itself or on human expert-curated interactions (e.g., no interactions are based on text-mining or associations). The dataset is curated to only include interactions derived from direct biophysical protein-protein interactions, metabolic pathway interactions, regulatory protein-DNA interactions, and kinase-substrate interactions. The details of the curation of these can interactomes can be found in (Zitnick et al., 2019).

The evolutionary history of the set of PPIs was obtained by (Zitnick et al., 2019) and is derived from a high-resolution phylogenetic tree (Hug et al., 2016). The tree is composed of Archaea, Bacteria, and Eukaryota and captures a diversity of species in each lineage. The phylogenetic tree is used to characterize the evolution of each species based on the total branch length (which takes the form of nucleotide substitutions per site) from the root of the tree to the leaf of the species. The phylogenetic taxonomy, the names of species, and lineages of each species were taken from the NCBI Taxonomy database (Federhen, 2011). Details of how this is associated with each species can be found at (http://snap.stanford.edu/tree-of-life), and we refer to (Zitnick et al., 2019) for further specifics on how each species was assigned an average nucleotide substitution rate. Ultimately these protein interactomes are incomplete models that may change as time goes on. Because we do not wish to bias our results, our statistical analyses were performed only over the interactomes of the species based on more than 1000 citations in the literature.

### Spectral analysis to find macroscales with EI^max^

Spectral methods have proved to be successful in identifying good graph partitions in a wide variety of applications (Guattery & Miller, 1995). Given an undirected network, we take the degree-normalized adjacency matrix *A* and compute the eigendecomposition *A* = *E*^*E*^*T*^, where the *i*^*th*^ column of *E* is the normalized eigenvector corresponding to the *i*^*th*^ eigenvalue, and *Λ* is the matrix with the *i*^*th*^ eigenvalue on the *i*^*th*^ diagonal and zeros elsewhere. The eigenvector matrix *E* contains rich information about the structure of the network, including information about the optimal scale of a network. The rows of *E* correspond to nodes in the network, so we construct a vector representation of each node’s contribution to the network topology by weighting the columns of *E* by their corresponding eigenvalues, removing columns that correspond to null eigenvalues, and associating the resulting row vectors with the nodes of the network. We construct a distance metric that reflects similarity in causal structure between pairs of nodes by taking the cosine similarity between the vectors corresponding to nodes. If a pair of nodes are not in each other’s Markov blankets, coarse graining them together cannot increase the effective information, so we define the distance between them to be ∞ (or simply very large, in this case, 1000). We use this metric to cluster the nodes of the network using the OPTICS algorithm (Ankerst et al., 1999) which we can interpret as a coarse-graining to construct a macroscale network, where micro-nodes are placed in the same macro-node if they are placed in the same cluster. Note that this method for detecting causal emergence in networks is explore in detail in other sources (Klein & Hoel, 2020).

### Robustness of causal emergence differences across species interactomes

To ensure that the differences observed in the causal emergence values of the protein-protein interaction networks were not merely a statistical artefact, we conducted a series of robustness tests of our analysis. These tests were necessary for two key reasons. First, the nature of interaction data in biology is inherently difficult to obtain. While many of the tools we use to collect, clean, and interpret biological systems are sophisticated, they are nonetheless subject to potential biases. However, if there were systematic biases in the network construction process for the protein interactomes used in this study (for example, if the interaction networks of eukaryotic species systematically over-estimated certain interactions), randomization procedures should clarify the extent to which the results we observed are truly a property of the species themselves.

Second, these robustness tests offer insights into whether there is anything intrinsic to the network structures of the eukaryotic or prokaryotic species that could be contributing to their causal emergence values. For example, the protein interaction networks of the eukaryote, *Rattus norvegicus* (the common sewer rat), has a certain amount of causal emergence. Would an arbitrary, simulated network with the same number of nodes and edges, connected randomly, also have a similar amount of causal emergence? By performing a series of robustness tests on the protein interaction networks in our study, we can get closer to the question of whether or not there is anything intrinsic to the protein interaction network of *Rattus norvegicus*, or any other species, that makes it particularly prone to displaying higher-scale informative structures?

To address the two concerns above, we performed two separate but similar robustness tests. The first uses a network null model known as the *configuration model* in order to randomize the connectivity of the protein interactomes while also preserving the number of nodes, edges, and distribution of node degree (Garlaschelli & Loffredo, 2008). The second robustness test involves random *edge rewiring* (Karrer, Levina, & Newman, 2007). For each network in our study, we iteratively increased the fraction of random edges to rewire in the network; an edge, *e*_*ij*_, that connects nodes *v*_*i*_ and *v*_*j*_, becomes re-connected to a new node, *v*_*k*_, forming a new edge, *e*_*ik*_, instead of the original *e*_*ij*_. We do this with an iteratively-increasing fraction of edges, starting with 1% of edges and increasing until 100% of the network’s edges are rewired.

If the causal emergence values of the networks in this study decrease following the robustness tests above—and in particular if they decrease *differently* for Eukaryota and prokaryotes—then the differences we observe are unlikely to have arisen simply from chance, noisy/biased data, or otherwise coincidental, *ad hoc* network properties. Instead, our testing of the robustness of our analysis lend credence to the main finding of this paper, which is that species that emerged later in evolutionary time are associated with more informative macroscale protein interaction networks.

In Fig. 4A, we show how the causal emergence of Archaea, Bacteria, and Eukaryota interactomes all decreases as a higher and higher amount of network edges are rewired, indicating that random rewiring has a similar effect on all datasets. This analysis suggests that if there were significant noise in the network data itself (i.e., connections between proteins where there otherwise should not be or a lack of connections where there should be), we should not expect to see the magnitude of causal emergence values that we indeed do see. This adds evidence that the inherent noise in the data collection process is not sufficient to produce the results we see.

In Fig. 4B, we show that random null models of the networks used in this study are characteristically unlikely to have values for causal emergence values that are at all similar to the original interactomes. On the contrary, the *maximum* average causal emergence value for any of the networks used here reaches only 3% of the original network’s values. This suggests that random null models of networks are less likely to contain higher scale structure but also that the observed differences in the causal emergence values for prokaryotic and eukaryotic species is unlikely to be driven merely due to basic properties like their edge density or degree distribution.

While it is impossible to exhaust all possible sources of bias or confounding variables in biological networks, the two statistical controls performed here get us closer to validating the hypotheses underlying this work: that evolution brings about higher informative scales in protein networks.

## Acknowledgements

M.L. gratefully acknowledges support by the Allen Discovery Center program through The Paul G. Allen Frontiers Group (12171). This publication was supported by a grant from Templeton World Charity Foundation, Inc. (TWCFG0273). The opinions expressed in this publication are those of the author(s) and do not necessarily reflect the views of Templeton World Charity Foundation, Inc. The project was also supported and made possible by the Army Research Office (proposal 77111-PH-II). B.K. acknowledges support from the National Defense Science and Engineering Grant (NDSEG).

## Author contributions

E.H., B.K., and M.L. conceived the project. E.H., B.K., A.S. and M.L. wrote the article. E.H., B.K., and R.G. performed the analyses.

